# Bacterial quorum sensing allows graded and bimodal cellular responses to variations in population density

**DOI:** 10.1101/850297

**Authors:** Jennifer B. Rattray, Stephen A. Thomas, Yifei Wang, Evgeniya Molotkova, James Gurney, John J. Varga, Sam P. Brown

## Abstract

Quorum sensing (QS) is a mechanism of cell–cell communication that connects gene expression to environmental conditions (e.g. density) in many bacterial species, mediated by diffusible signal molecules. Current functional studies focus on a dichotomy of QS on/off (or, quorate / sub-quorate) states, overlooking the potential for intermediate, graded responses to shifts in the environment. Here, we track QS regulated protease (*lasB*) expression and show that *Pseudomonas aeruginosa* can deliver a graded behavioral response to fine-scale variation in population density, on both the population and single-cell scales. On the population scale, we see a graded response to variation in environmental population density. On the single-cell scale, we see significant bimodality at higher densities, with separate OFF and ON sub-populations that respond differentially to changes in density; static OFF cells and increasing intensity of expression among ON cells. Together these results indicate that QS can tune gene expression to graded environmental change, with no critical cell mass or ‘quorum’ at which behavioral responses are activated on either the individual cell or population scale. In an infection context, our results indicate there is not a hard threshold separating sub-quorate ‘stealth’ mode and a quorate ‘attack’ mode.

## Introduction

Many species of bacteria are capable of a form of cell-cell communication via diffusible signal molecules, generally referred to as quorum sensing (QS). The study of QS has largely focused on the intracellular gene regulatory scale, leading to a detailed understanding of the regulatory mechanisms shaping the production of and response to signal molecules in model organisms such as *Vibrio cholerae, Bacillus cereus* and *Pseudomonas aeruginosa* [1–3]. We now understand that QS is mediated by multiple diffusible signals that together control a diverse array of responses, including swarming, luminescence, competence and the production of diverse secreted factors [4,5]

While the molecular mechanisms of QS have been described for model organisms in remarkable detail, the functional and evolutionary context of QS continues to be disputed. In other words, while we now have a better understanding of *how* QS works, we still have limited understanding of *why* bacteria use this system to control behavior. What are the functions of QS? How do these QS functions help bacteria to survive and grow? The standard answer is that bacteria use QS to sense when they are at sufficient density (‘quorate’) to efficiently turn on cooperative behaviors such as secretion of toxins and enzymes in order to collectively modify their environment [6].

Other researchers have argued that QS is an asocial sensing apparatus, where individual cells produce and monitor signal levels in order to infer their physical environment (am I in an open or enclosed space?) [7]. More recently, integration of molecular and evolutionary approaches has increased the menu of potential functions to include sensing multiple aspects of both the social and physical environment [6,8–10] and coordinating complex social strategies that limit the profitability of non-cooperating ‘cheat’ strains [11–18].

A critical step in assessing the various adaptive hypotheses is establishing the functional capacities and limits of QS. Previous studies have demonstrated ‘density sensing’ functions – populations can use QS to sense when they exceed a density threshold [6,19,20]. In addition, Darch et al. (2012) demonstrated that responding with increased QS controlled cooperative activity at high density can provide a fitness benefit [6]. Other studies have demonstrated ‘diffusion sensing’ functions [7] – QS systems can functionally respond to variation in physical containment, so that even a single cell can become ‘quorate’ (turn on a QS controlled reporter gene) if isolated in a sufficiently small contained space [9]. More recently, some studies have demonstrated ‘genotype sensing’ functions – QS can respond to variation in the genotypic composition of a population, restricting QS-controlled responses to populations that are enriched with wildtypes [11,14,21,22].

The functional studies outlined above largely focus on a dichotomy of QS on/off (or, quorate / sub-quorate) states, overlooking the potential for intermediate, graded responses (Fig 1A). The threshold quorate/non-quorate concept is ingrained in the QS literature following the use of the legal ‘quorum’ analogy [20], and is also supported by mathematical models of QS signal dynamics that highlight how sufficiently strong positive feedback control of signal production can produce a sharp threshold response to changes in environmental parameters such as density or diffusion [23,24]. However, these same mathematical models indicate that graded responses are also possible, dependent on the model parameterization. More generally, Fig 1A highlights that the phenotypic response of QS bacteria to differing environmental conditions can be viewed as a ‘reaction norm’ [25–28] that can in principle take differing shapes. Reaction norms describe phenotypic responses of a single genotype (y-axis, Fig 1A) to varying environmental inputs (x-axis, Fig 1A). Incorporating a reaction norm framework provides a menu of quantitative metrics to define QS responses to environmental variation (e.g., slope, intercept, and variances). With this reaction norm framework, it is important to emphasize that in our study the x-axis is not time, but instead captures a gradient of environmental conditions. Whether responses are graded or thresholded during the growth towards high density is a separate line of inquiry [29]. Describing the reaction-norms of QS cells and populations to contrasting environments is an important step towards understanding the capacities of QS systems to differentially respond to novel environments.

**Figure 1.**
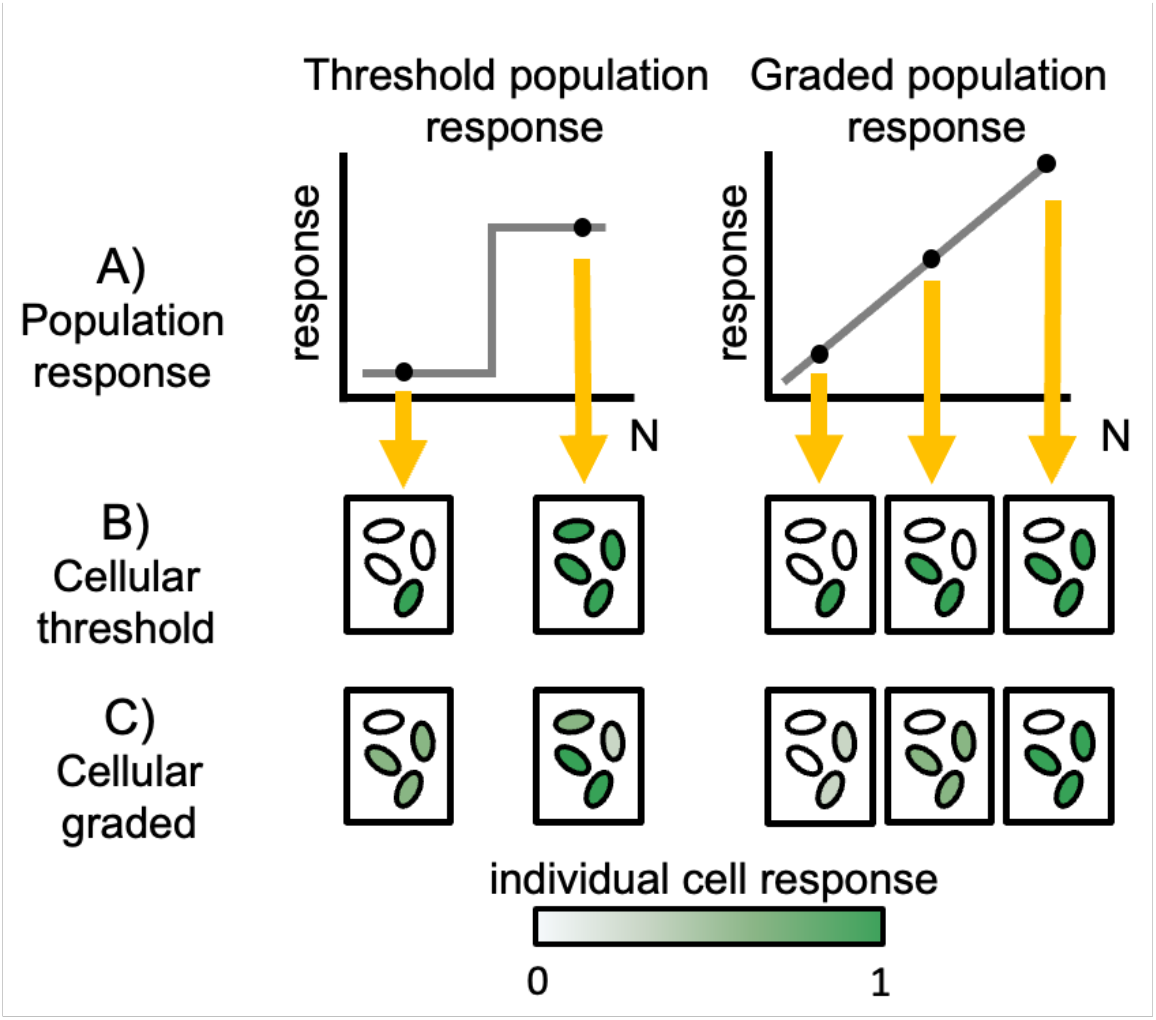
Schematic of potential population and single cell responses to variation in cell density. **A**) Population response (y-axis) across discrete carrying capacity environments (N, x-axis), given a threshold (left) or graded response (right). In (B) and (C) we outline alternative cell-scale responses (intensity of green cells) that are consistent with discrete population scale behaviors (yellow arrows). (B) threshold (ON/OFF) cellular responses can produce a threshold or graded responses on population scale. (C) graded individual responses can produce threshold or graded responses on a population scale.

Whether the population scale reaction norm to environmental variation is threshold-like or graded (Fig 1A), a separate question is how collective population-level responses are constructed out of individual cellular contributions (Fig 1B,C). Studies of QS on a single-cell scale have revealed substantial heterogeneity in response to QS signals [9,30–36], highlighting that cell-cell communication does not necessarily result in tight synchronization of individual cell activity (Fig 1B,C). In some systems, heterogeneity can be quenched by the addition of extra signal [31,33], implying a lack of receptor saturation. However, this is not a universal result [30], indicating that other molecular processes can drive cellular variation in response. Regardless of the molecular details, we currently lack a behavioral understanding of how individual cellular responses vary with changes in the environment.

In the current study we address the canonical ‘density sensing’ function of QS, using the environmental generalist and opportunistic pathogen *Pseudomonas aeruginosa*, and an unprecedented scale of environmental resolution (13 discrete limiting carbon levels conducted in triplicate, generating 39 density environments). QS in *Pseudomonas aeruginosa* is heavily studied in a high-density context, revealing a complex mechanism of multi-signal control [37–41]. Our first challenge is to map the population scale resolving power of QS to quantitatively discriminate graded differences in population density (Fig 1A). Does *P. aeruginosa* respond in a purely threshold manner, collapsing quantitative differences in population density into a simple low / high qualitative output, or can QS allow *P. aeruginosa* to deliver a graded response to distinct environmental densities? Our second challenge is to understand how collective responses are partitioned across individual cells. Are changes in collective responses governed primarily by changes in the proportion of cells in an on state (Fig 1B) or changes in the individual cell intensity of response (Fig 1C), or both?

## Results

### Collective level of response to density is graded and linear

Our first challenge is to map out the population scale reaction norm of the collective QS-controlled protease (*lasB*) response to variation in population densities. To provide a detailed picture of the QS response reaction norm to varying density, we grew a QS reporter strain (PAO1 pMHLAS containing the *PlasB::gfp(ASV)* reporter construct for QS regulated protease expression [42]) under 13 conditions of carbon limitation in triplicate and measured average fluorescence output per cell as the populations reach carrying capacity (Fig 2). Dead cells with compromised membranes were identified with a propidium iodide stain and excluded from analysis. The range of cell densities generated from this method is from 1×10^8^ cells/ml to 2×10^9^ cells/ml. Figure 2 shows that QS response is linear with increasing culture density, providing intermediate levels of average per-capita response to intermediate densities. To confirm the lack of threshold behavior we assessed alternate statistical models including threshold functions, and found that a linear fit model supports the data better than a step-function fit (AIC linear: 89, AIC step-function: 190; relative likelihood that the linear model is the best fit compared to step-function > 10^9^, see [43]), supporting a graded population response as outlined in Figure 1. This agrees with literature that QS induction at lower population densities is possible [6,9,19], but differs in that there is no observable population density at which populations ‘switch’, or reach quorum, into a responsive state.

**Figure 2.**
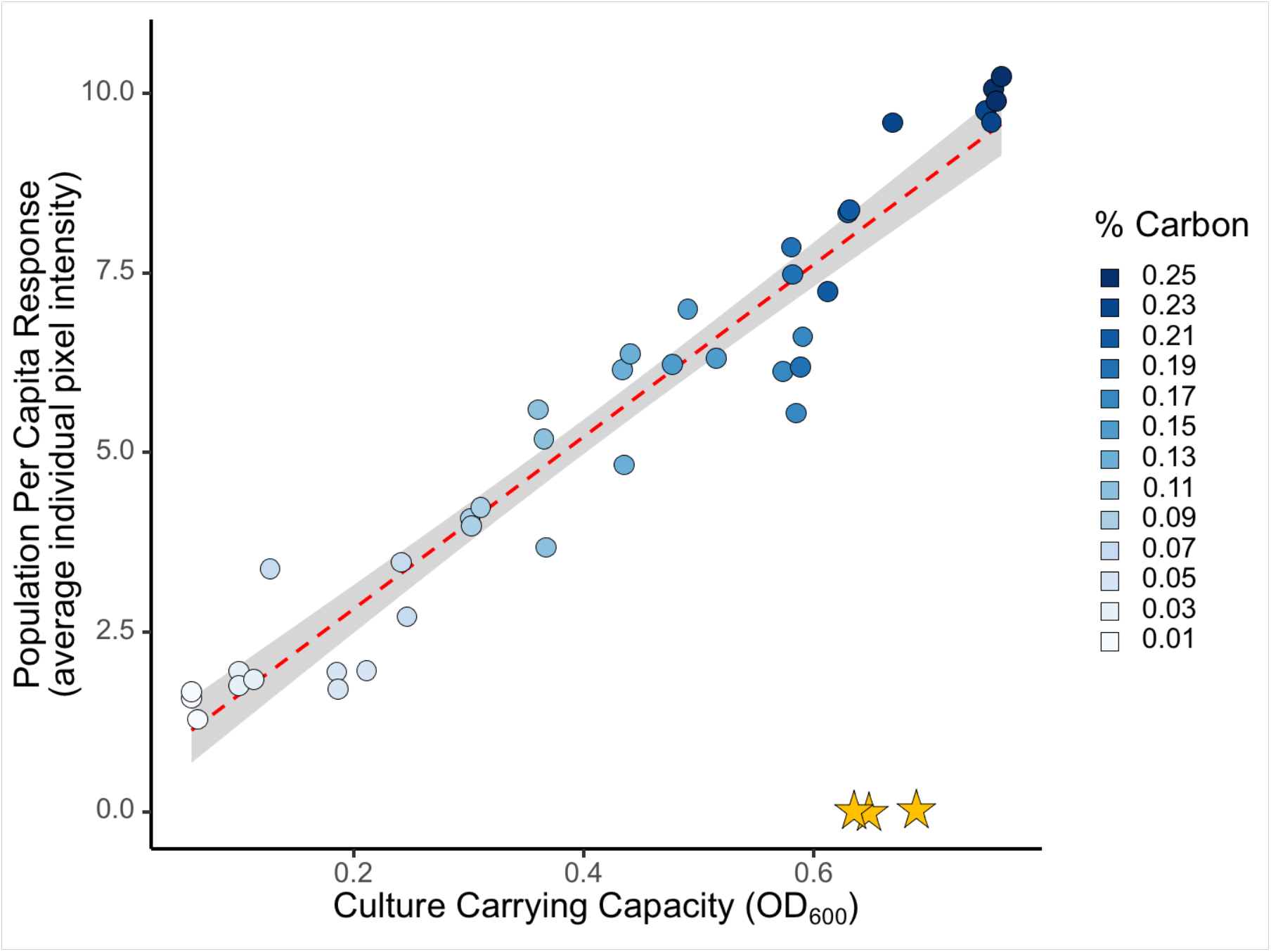
Population response to increasing cell density is linear and graded. 13 distinct culture carrying capacities were generated by manipulating the concentration of casein digest as the limiting resource (Fig S1). Cells were grown to carrying capacity in triplicate and immediately assayed for QS response via fluorescence microscopy imaging. Response is determined by a fusion of the quorum sensing controlled *lasB* promoter and an unstable green fluorescent protein (PAO1 pMHLAS containing *PlasB::gfp(ASV))*. Individual cell pixel intensity is a measure of cellular quorum sensing response and average pixel intensity is calculated across all cells in the population as a proxy for total population expression. Microscopy averages are congruent with population scale plate reader results (Figure S2). A quorum sensing signal knockout (Δ*lasI*Δ*rhlI*), yellow star, shows background response with no signal in the environment. Average population investment in QS increases as culture density increases with no observable density threshold (AIC linear: 89, AIC step-function: 190).

### Individual response to density is bimodal at high densities

Figure 2 establishes that on a collective population scale, the response to environmental variation (in density) is smoothly graded. Next, we ask how this collective response is built from individual cell contributions. Is the graded increase due to more cells turning on at higher densities (Figure 1B), cells turning on to a greater extent (Figure 1C), or both? To address this question, we take the same data presented in Figure 2 and now present the distribution of individual cellular responses rather than simply the mean response (Figure 3).

**Figure 3.**
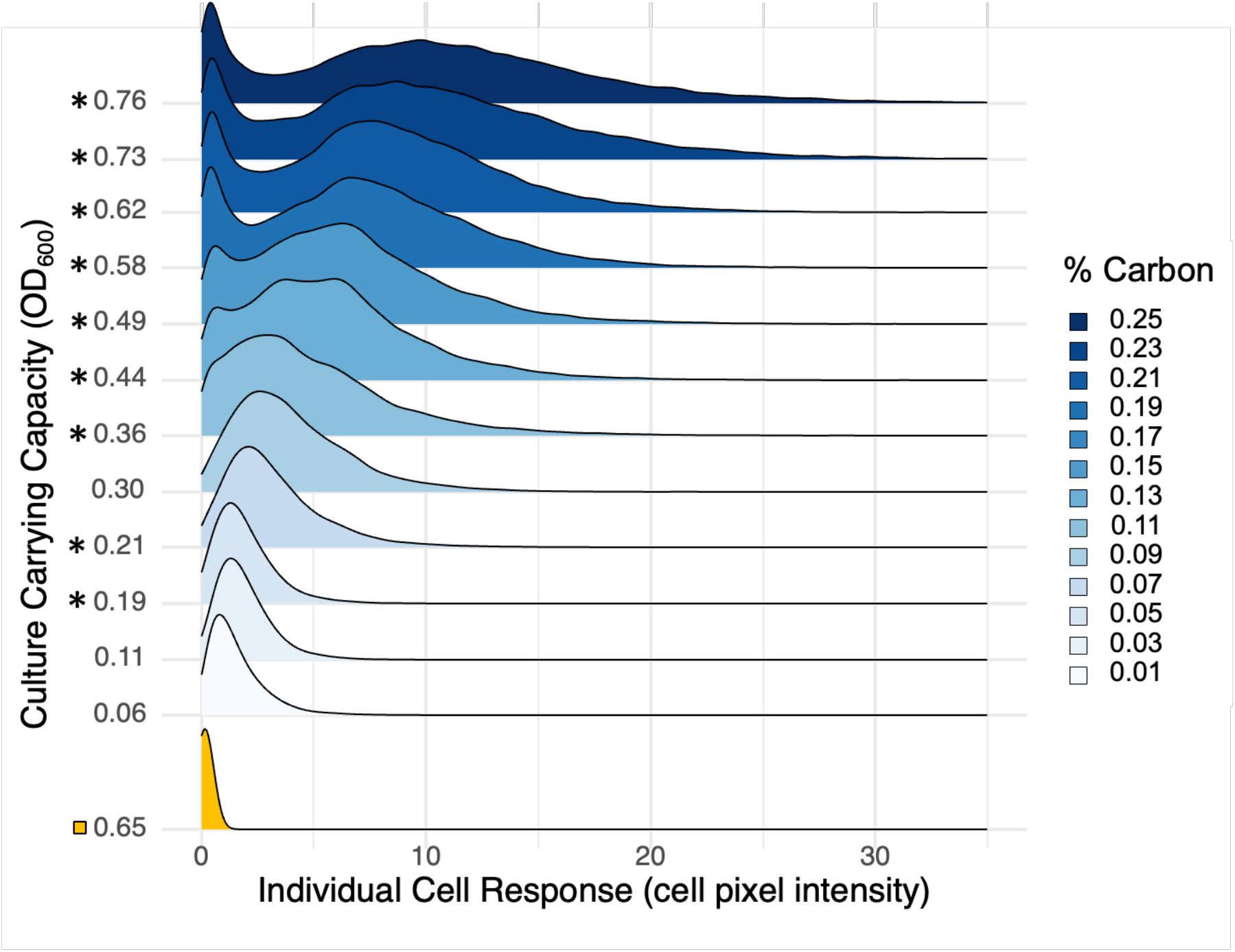
Individual response is heterogenous and bimodal at higher densities. Ridgeline density plot (bandwidth = 0.435) of single-cell *lasB* reporter response data showing the distribution of individual cell QS expression across the population. For brevity and plotting purposes, carrying capacities were averaged across 3 replicates for each of the 13 carbon environments before plotting. A full plot of each independent replicate environment can be found in Figure S3. Each line summarizes 18,000 to 30,000 individual cell measurements, scaled to a unit height. Asterisks indicate significant bimodality (Hartingan’s Dip Test [44], Figure S4). The quorum sensing signal knockout (*ΔlasIΔrhlI*) is denoted with yellow boxes. A total of 345,000 individual cell measurements were analyzed.

As expected from prior studies in other QS systems [9,30–36], plotting all individual responses within a population shows cell-to-cell variation in QS response within a single population despite isogenic and homogenous culture conditions (Figure 3). In addition, at higher densities we see significant bimodality (defined by Hartigan’s Dip Test, Figure S4), with the population segregating into an unresponsive, sub-quorate, OFF state and a responsive, quorate, ON state.

In light of this bimodality, we fit a two-component finite mixture model to the data (Figure 4A, see Figures S10-S13 and Table S2 for extended analysis), which allows us to define the average intensity of the ON state (Figure 4B) and the proportion of cells in the OFF or ON states (Figure 4C).

**Figure 4.**
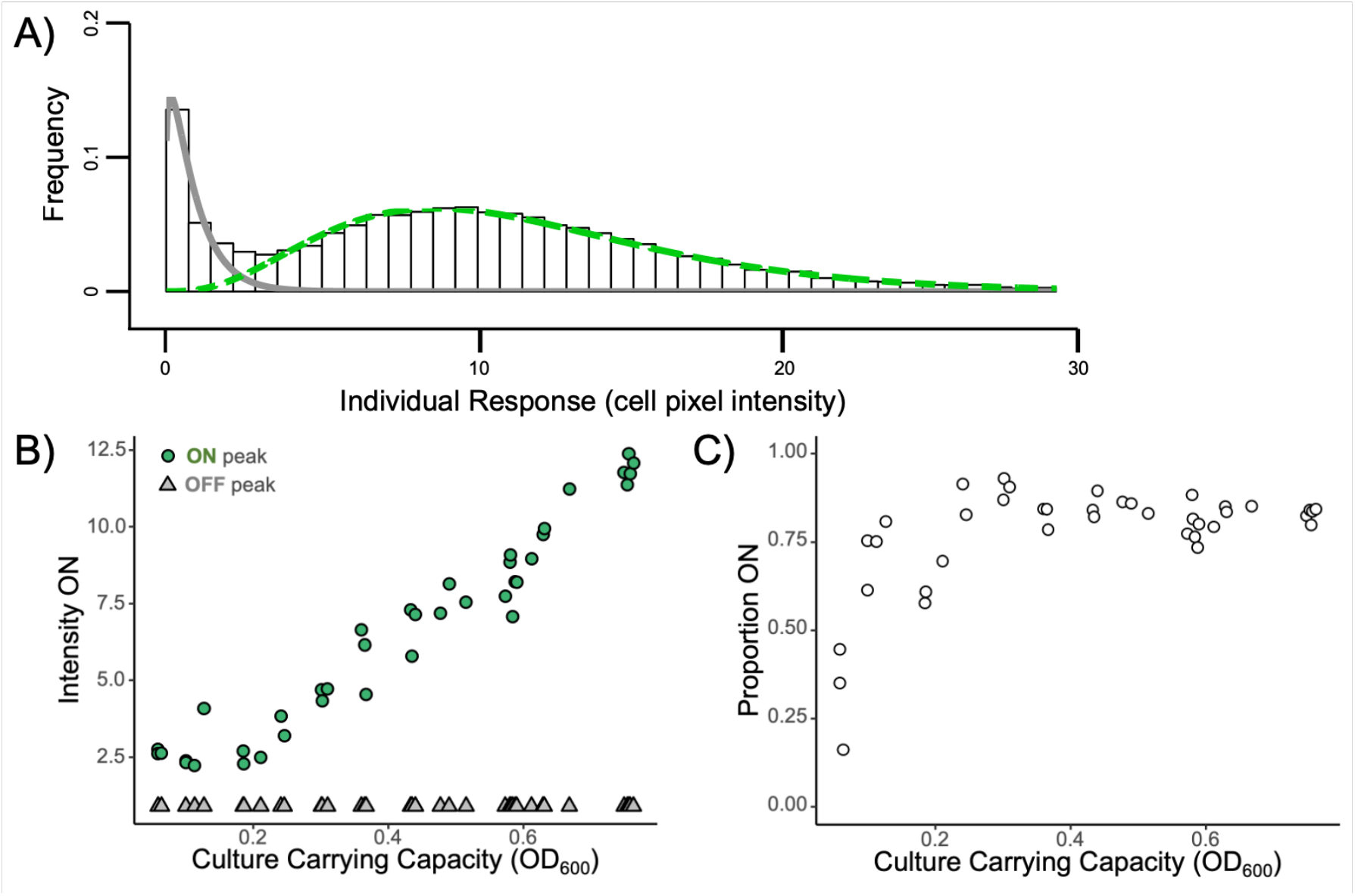
Proportion of cells responding and level of response varies with density. In light of the bimodal responses in Figure 3, we course-grain the single-cell *lasB* response data into discrete ON/OFF states. A) Method summary. We quantify distinct ON/OFF states by fitting a two-component finite mixture model at each measured optical density, where the OFF state is fixed to the OFF state of the highest density environment. The histogram shows the distribution of cellular expression levels at a single density treatment (0.76 OD_600_), the grey line is the fitted OFF state and the green dashed line is the fitted ON state. B) The mean intensity of the ON (green circle) and OFF (grey triangle) states is determined from the means of mixture model component fits (green and grey lines in panel A). The mean intensity of the ON state distribution increases as culture density increases, while the mean of the OFF state remains constant. C) The proportion of cells ON in the population is determined from the relative mass of cells in the model component fits. The proportion ON increases with culture density but does not reach 100%.

Figure 4B illustrates a graded linear increase in the intensity of the ON state with increasing environmental density, and a density-invariant off state. Figure 4C illustrates that the proportion of cells that are ON plateaus at around 85% at densities with consistent support for bimodality (above 0.36 OD_600_). At lower densities, the intensity of the ON state (Figure 4B) declines to a point where the OFF and ON states are no longer significantly different and the dip test fails to reject uni-modality (Figure S4). In supplemental materials, we present alternate statistical analyses of this data, and of other related datasets. Across other experiments, we find consistent support for the graded and bimodal response pattern on the single-cell scale across multiple assay time-points (Figure S5) and across two reporter strain constructs (Figure S6) and support for the graded and linear response pattern on the population scale across fluorescent and lux reporters (Figure S2, Figure S7). We find further support for the graded reaction norm on the population scale across two additional QS controlled genes (*pqsA, rhlI;* Figure S7).

## Discussion

Our results show that populations of *P. aeruginosa* can respond in a smoothly graded manner to variation in environmental density (Figure 2), that populations exhibit significant bimodality at higher densities (Figure 3), and that this population scale graded response can be described by the number of responsive ‘ON’ cells and the intensity of the ‘ON’ state (Figure 4). The ability to achieve a graded population scale response implies in principle that *P. aeruginosa* can tune collective responses (such as the secreted elastase virulence factor produced by our *lasB* reporter gene) to graded environmental changes, rather than simply course-graining into a simple ‘high / low’ dichotomy. A similar population scale graded response to continuous environmental variation is visible in the data from Allen et al., which looked at variation in the genotypic composition of mixed populations grown to the same total density [11]. As the proportion of wildtype (PAO1 versus Δ*lasR* ‘cheats’) increased, the wildtype per-capita investment in cooperative LasB secretions also increased, providing a simple behavioral mechanism to protect cooperative investments from exploitation by cheats [11,22].

The existence of graded population scale responses across two continuously varying environmental inputs (density, genotypic composition) raises the question of why use a graded response? Is there an evolutionary rationale for a graded response, or is a graded increase simply the ‘best approximation’ of a threshold response, given a simple system working under genetic constraints? Existing evolutionary theory suggest that graded investment reaction norms can be adaptive, under a range of distinct scenarios [45,46] (34, 35). In the specific context of quorum-sensing bacteria, evolutionary theory suggests that population scale responses to increasing density should depend critically on the shape of the cost and benefit functions of increasing cooperative investments. Specifically, a graded response is predicted to be the optimal strategy if the benefit function is decelerating and costs are linear with increasing investment [47].

To further consider the functional context of the graded reaction norms, we turn to the single cell scale data, which reveals how the graded population response is built from the contributions of individual cells. In agreement with previous work in multiple quorum-sensing organisms [9,30– 36,48,49], we find cell-scale heterogeneity. In addition, our results illustrate how cellular heterogeneity changes with the environment, demonstrating the onset of ON/OFF bimodality at intermediate densities, with both the proportion of cells ON and the intensity of the cellular ON states increasing with increases in culture carrying capacity (Figures 3 & 4).

The presence of a bimodal QS response is in contrast with the common view of QS as a mechanism of cell synchronization yet can be viewed as a striking example of widely observed cellular heterogeneity under QS control. Indeed, bimodal responses are implicit in some of the previous single-cell QS literature [31,32,49], for example Darch et al. (2018) report distinct populations of QS-responsive and non-responsive cells within single experimental runs [49]. The degree of heterogeneity in any cellular trait can be interpreted as the interplay of biochemical properties of molecules and the architecture of gene-regulatory networks [50]. Given that regulatory networks are subject to mutation and selection, this implies that the degree of heterogeneity is an evolvable trait [51]. In the context of QS, positive feedback loops (signal auto-regulation [52]) and the presence of cooperative transcription factor binding [53] provides recognized regulatory ingredients for bimodal expression [54]. Recently, the presence of heterogeneous QS response at the single-cell scale has been ascribed to a potential bet-hedge against mis-directed QS induction [48], suggesting that our OFF cells are poised to more quickly resume growth in the event of a rapid return to a growth-friendly environment.

We made a number of specific observational choices in order to conduct our experiment that could have shaped our results in ways that are not generalizable to other contexts. In the supplementary we detail a number of additional experiments (and alternate statistical analysis approaches) that collectively illustrate the robustness of our findings. In brief, we found that our single cell results are not sensitive to the time the population was sampled (Figure S5), the presence of a potentially leaky P*lac::lasR* on the pMHLAS construct (Figure S6), or the plasmid nature of the pMHLAS construct (Figure S6). Additionally, we recognize that *lasB* is only one gene out of hundreds that are controlled by QS [3], and is often co-regulated by other factors [55– 57]. We chose to initially focus on *lasB* as it is a traditionally studied QS-controlled trait [58–60], it is under multi-signal control [10,61] and has clinical significance as a virulence factor [62,63]. To begin to address the generality of our results across genes in *P. aeruginosa*, we show that two other QS regulated genes with complex promoters, *pqsA* and *rhlI*, also support a graded population response (Figure S7). It remains to be seen whether the graded responses we report here are consistent across all QS controlled genes in *P. aeruginosa*, and across QS systems in other species,

A recent transcriptomic analysis of clinical versus *in vitro* gene expression in *P. aeruginosa* called into question the clinical relevance of *in vitro* models of QS, reporting that QS activity (including *lasB* expression) was systematically higher in *in vitro* models [64]. Our results provide a simple interpretation of this difference: *in vitro* models are typically conducted under higher experimental densities, resulting in higher levels of average QS gene expression (Figure 2). Consistent with this graded response interpretation, Cornforth et al. (2018) also reported higher levels of relative expression in *in vitro* biofilm models (close-packed cells, the highest local density achievable) compared to *in vitro* planktonic models.

In summary, our results provide a finely resolved mapping of the QS reaction norm to environmental density in PAO1, on both the collective and single-cell scale. On the population scale we see a graded linear response across a range of cellular densities (1×10^8^ cells/ml to 2×10^9^ cells/ml) and significant individual-scale bimodality at higher densities. We further resolve this linear population response (Figure 2) into a combination of the likelihood of being responsive and the intensity of response (Figure 4). In an infection context, our results indicate that there is no hard threshold separating sub-quorate ‘stealth’ mode and a quorate ‘attack’ mode [65]. One implication is that attempts to control virulence and biofilm expression in medicine and industry via QS inhibition could have impacts across a wider spectrum of population densities. In this applied context, it is important to assess the generality of our results and ask, how do QS reaction-norms vary across strains and species of QS bacteria? How do they vary across environments? More broadly, our work undermines the threshold concept of a ‘quorum’, instead placing QS bacteria in the graded world of reaction norms.

## Materials and Methods

### Bacterial Strains and Growth Conditions

The two main bacterial strains used in this study are *P. aeruginosa* NPAO1 (Nottingham-PAO1) containing the *PlasB::gfp(ASV)* quorum sensing reporter pMHLAS [42] and a double signal synthase mutant incapable of producing QS signal molecules, *P. aeruginosa* NPAO1 *ΔlasI/ΔrhlI* containing the same *PlasB::gfp(ASV)* quorum sensing reporter pMHLAS. A complete table of strains used in the main text and supplemental figures can be found in Supplemental Table 1. Overnight cultures were grown in lysogeny broth (LB), supplemented with 50 ug/ml gentamicin to maintain the pMHLAS plasmid, with shaking at 37 °C. Experiments were conducted in lightly buffered (50 mM MOPS) M9 minimal defined media composed of an autoclaved basal salts solution (Na_2_HPO_4_, 6.8 gL^−1^; KH_2_PO_4_, 3.0 gL^−1^; NaCl, 0.5 gL^−1^), and filter-sterilized 1 mM MgSO_4_, 100 uM CaCl_2_, and 1X Hutner’s Trace Elements with casein digest, as the sole carbon source (ThermoFisher Difco™ Casein Digest CAT 211610).

### Controlling Culture Carrying Capacity

We manipulated density by controlling the limiting resource in the media, carbon, allowing us to tune the carrying capacity of each treatment (Figure S1). To cover a variety of densities, we generated a carbon range between 0.05% and 0.25% via dilutions of a 0.5% carbon minimal media stock for a total of 13 different carrying capacities with three replicates each. This produced a range of densities environments from 1.18×10^8^ cells/ml to 2.02×10^9^ cells/ml. Overnight cultures were grown in LB gentamicin 50 ug/ml and centrifuged at 8,500 x g for 2 minutes. The cells were then washed twice with carbonless minimal media and then each carbon treatment was adjusted to OD_600_ = 0.05. Then, 200 uL of each sample was added to a 96-well microplate. Plates were incubated with shaking at 37 °C in a Cytation/BioSpa plate reader and growth curves were generated by absorbance (OD_600_) readings taken at 30-min intervals.

### Measuring Population QS Response

To measure population response, we performed growth-curve experiments as previously described using PAO1 *PlasB::gfp(ASV)*, additionally taking fluorescence readings at 30-min intervals. Fluorescence, population level response, was recorded when populations reached the end of their exponential growth phase, before they entered stationary phase. Background fluorescence of the reporter was determined with the QS signal deficient mutant PAO1 *ΔlasI/ΔrhlI PlasB::gfp(ASV)*. The population microplate data (Figure S2) and averaged microscope data (Figure 2) agreed, so the latter is provided in the primary text.

### Measuring Individual QS Response

To measure individual response, we performed growth-curve experiments as previously described, but removed samples for microscopy once cells reached end exponential phase. Since we control carrying capacity with the amount of carbon, the exact time that cells reach the end of exponential growth differs across treatments by 2-3 hours. To robustly sample cultures at this specific point, the slope of the two most recent time points on the growth curve was monitored and samples were taken as the slope approached 0. Replicate wells were kept growing to confirm that the treatment entered stationary phase right after the sampling time point. We also determined that our results are generalizable even when sampling at a pre-determined hour across concentrations (Figure S5). Samples were stained with propidium iodide to differentiate between life and dead cells and a small aliquot (5 ul) was added to a 0.01% poly-l-lysine coated slide to immobilize cells and immediately imaged to avoid changes in expression between sample acquisition and imaging in the dark on a Nikon Eclipse TI inverted microscope at 20x magnification. Live cell fluorescence microscopy was used for this study as fluorophores can be sensitive to fixation/permeabilization. These techniques can result in a decrease in fluorescence and therefore decrease in the observable dynamic range. Bright field, green fluorescence (20% Lumencor light engine power, 200ms exposure, and 64x gain-sufficient for imaging of low fluorescent cells without saturating pixel intensity), and red fluorescence (20% Lumencor light engine power, 800ms exposure, and 64x gain) channels were captured. Between 5,000 and 15,000 individual cells were captured for each sample. Aliquots were diluted immediately before imaging with carbonless minimal media when required to ensure an even distribution of cells.

### Single cell image analysis

A custom macro in ImageJ was written to analyze the image data, outlined in Figure S8. The macro uses ImageJ’s “analyze particles” command to identify single cells on the bright field image. This then generates a ROI (region of interest) for each individual cell and these ROIs were then overlaid onto the corresponding fluorescent image. The red fluorescence channel was used to identify dead cells with compromised membranes, which were excluded from further analysis. The green fluorescence channel reflected the QS reporter and pixel intensity was measured as a proxy for level of QS response. This tabulated live cell expression data was then analyzed using *Stata Statistical Software: Release 17* from StataCorp LLC. In order to improve the fit of the mixed models, the lowest pixel intensity measurement in the highest carbon PAO1 *ΔlasI/ΔrhlI PlasB::gfp(ASV)* treatment was subtracted from all pixel intensities so that expression started at 0.

### Statistical analysis summary

The analysis was done using *Stata Statistical Software: Release 17* from StataCorp LLC and the additional third party resources: [66–70]. Each of the 39 populations was fit to a finite mixture model of two Gamma distributions. The latent classes in the mixture model correspond to OFF and ON cells. Gamma distributions are preferred to Normal distributions as gene expression is strictly non-negative and necessarily right-skewed. The models provide maximum likelihood estimates of the proportion of cells in each latent class and the shape and scale parameters of the component Gamma distributions. Mean expression level for each distribution is the product of shape and scale parameters. Information criteria for aggregate mean expression level was also calculated using Stata.

## Acknowledgments

We thank Steve Diggle, Marvin Whitely, Kathleen O’Connor and members of the Center for Microbial Dynamics and Infection (CMDI) for valuable comments and discussion on this work. We thank Dr. Kasper Norskov Kragh for the *PlasB::gfp(ASV)* reporter construct. This research was supported by the National Science Foundation Graduate Research Fellowship Program under Grant No. DGE-1650044, The Simons Foundation Grant No. 396001, and National Institute of Health Grants 1R21AI143296 and 1R21AI156817.

## Supplementary Information for

### Data Supplement

**Figure S1.**
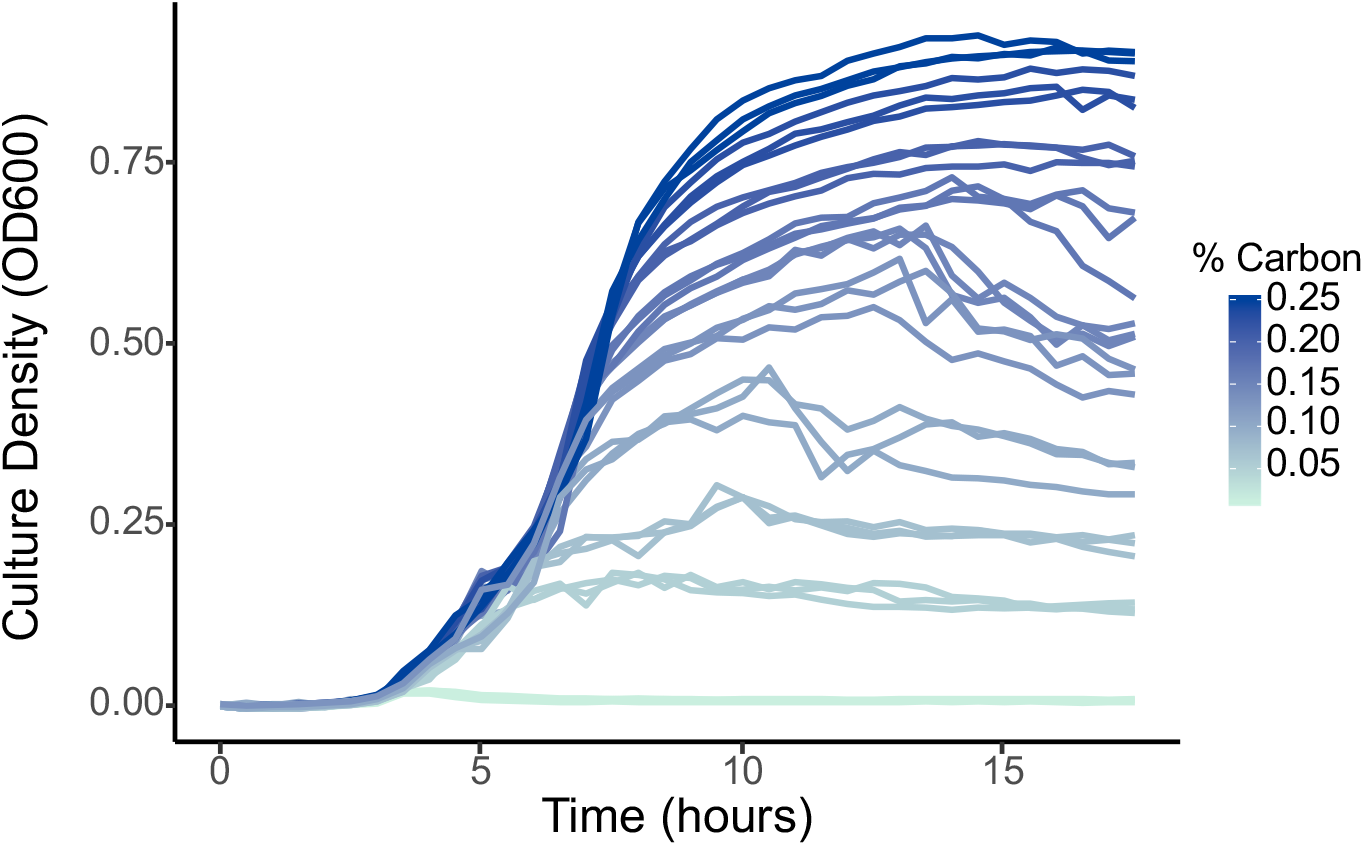
Growth curves for PAO1 pMHLAS across different carbon limiting environments. Discrete environmental densities can be generated by varying carbon availability, therefore manipulating the carrying capacity of the culture.

**Figure S2.**
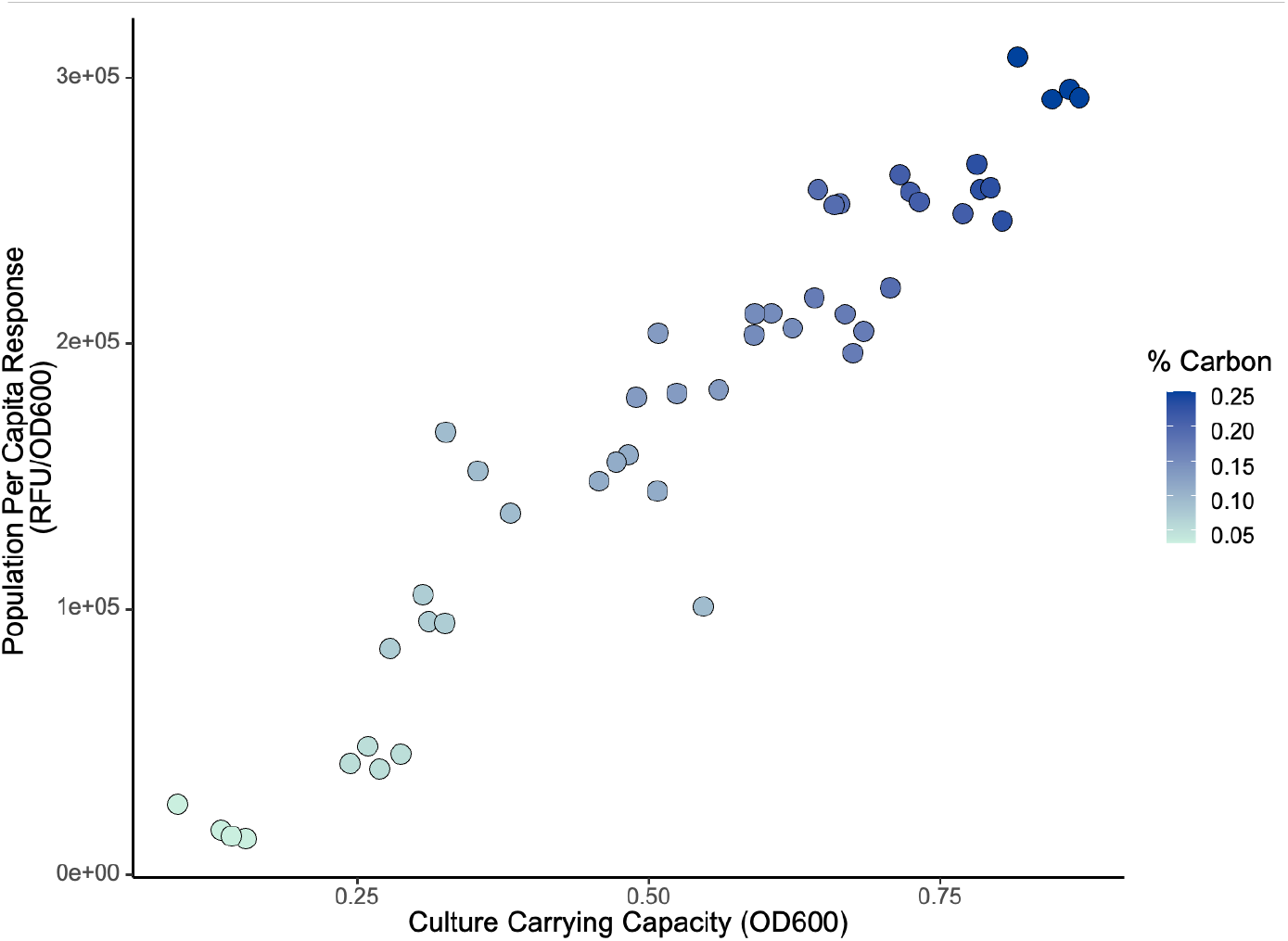
Microplate population per capita data is graded and linear. PAO1 pMHLAS was grown as mentioned in the methods and fluorescence and OD_600_ were measured on a Cytation Sense plate reader. Microplate results agree with microscopy results that that population response to increasing cell density is linear and graded. OD_600_ of 0.25 is 5.5×10^8^ cells/ml, OD_600_ of 0.50 is 1.3×10^9^ cells/ml, and OD_600_ of 0.75 is 2.05×10^9^ cells/ml.

**Figure S3.**
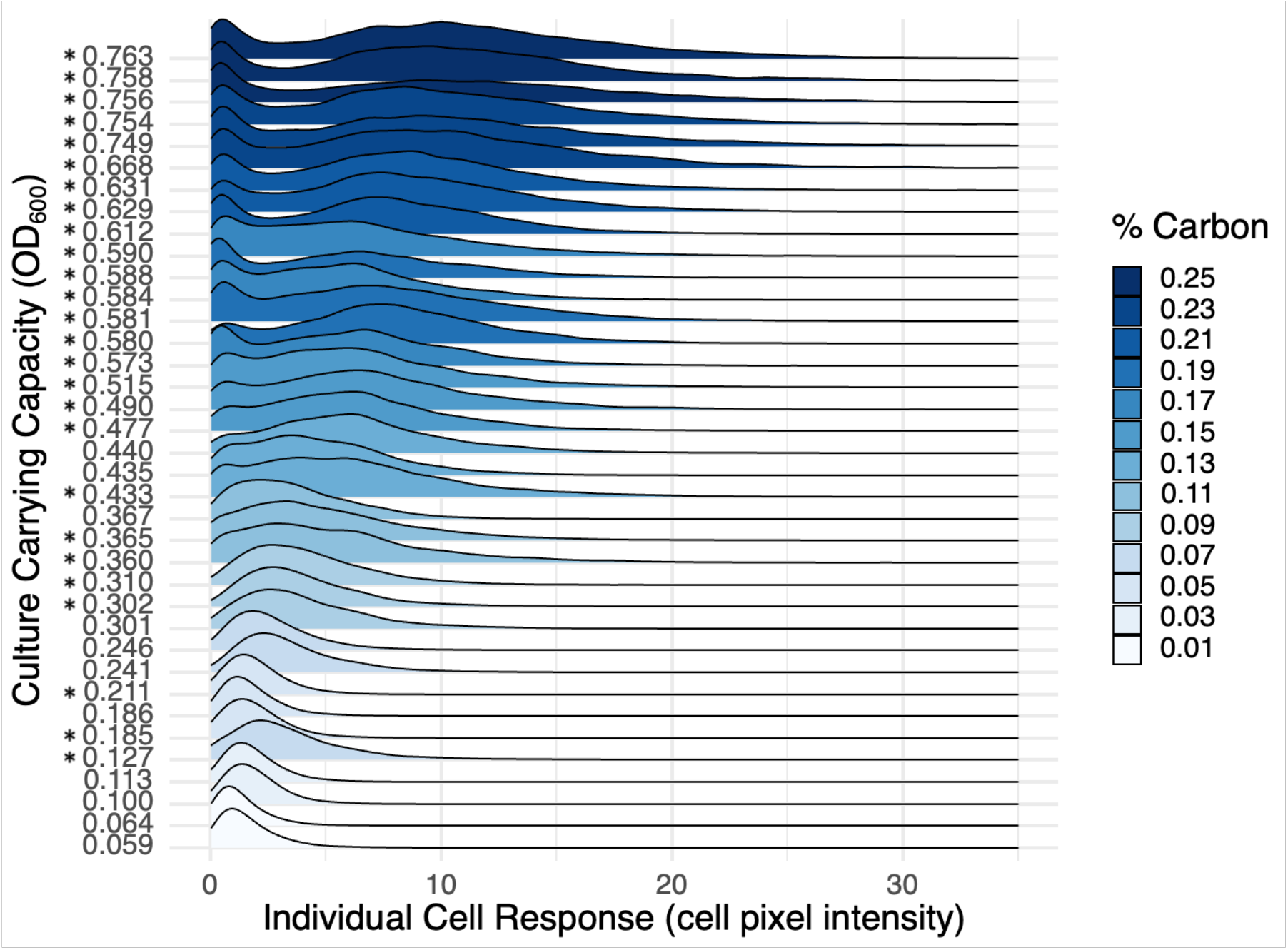
Ridgeline density plot (bandwidth = 0.529) of single-cell *lasB* reporter response data showing the distribution of individual cell QS expression across the population. All 39 replicates are plotted separately. Asterisks indicate significant bimodality (Hartingan’s Dip Test [44], Figure S4).

**Figure S4.**
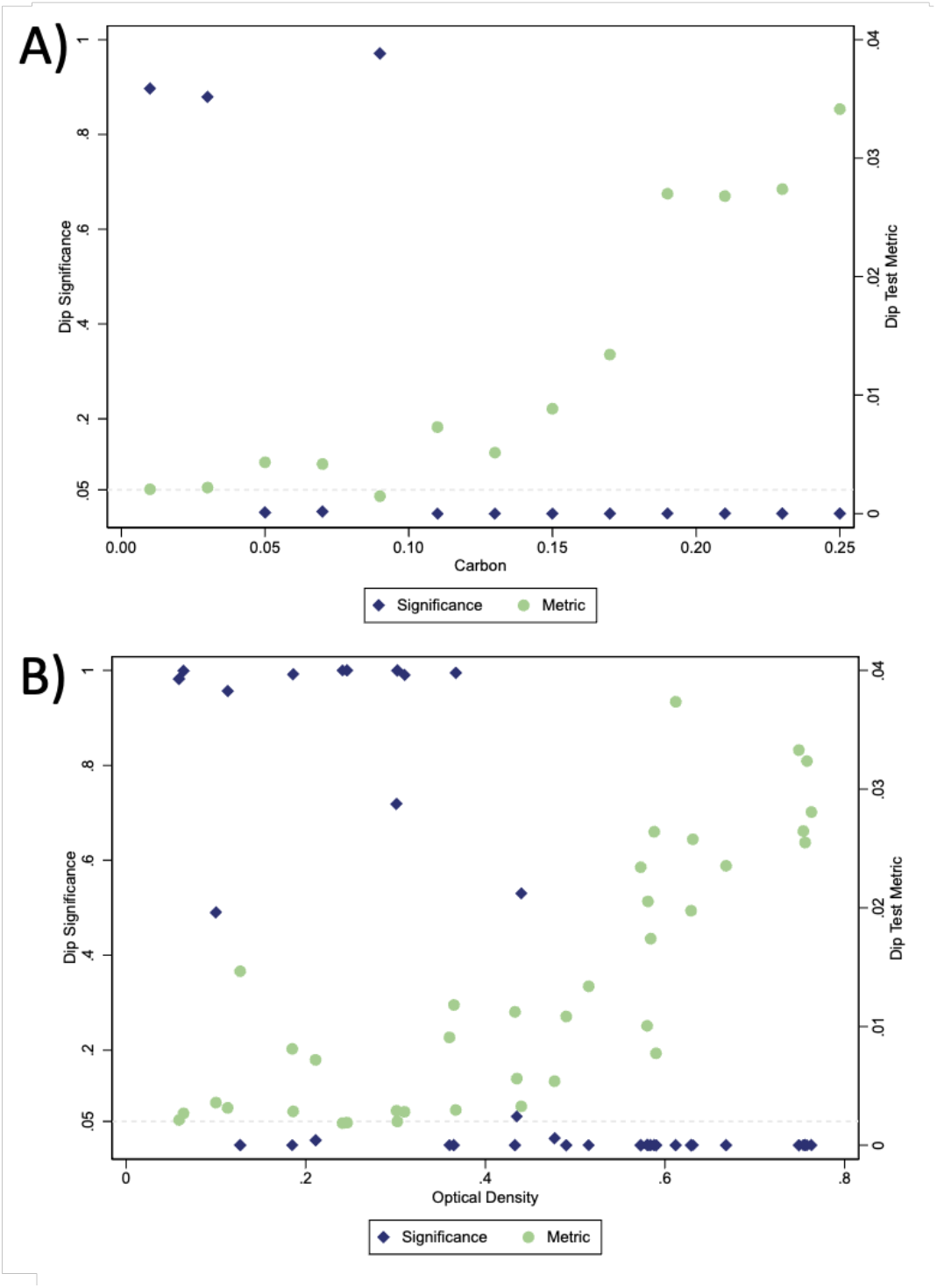
Hartigan’s Dip Test. A) Results of Hartigan’s dip test [44] for assessing bimodality of the expression level distributions at each carbon level for data shown in Figure 4. B) Results of Hartigan’s dip test for assessing bimodality of the expression level distributions at each OD level for data shown in Figure S3. Statistical significance in the form of p-value is plotted in yellow on the left axis, and the dip metric is plotted in green on the right axis. The most conservative p-value is shown.

**Figure S5.**
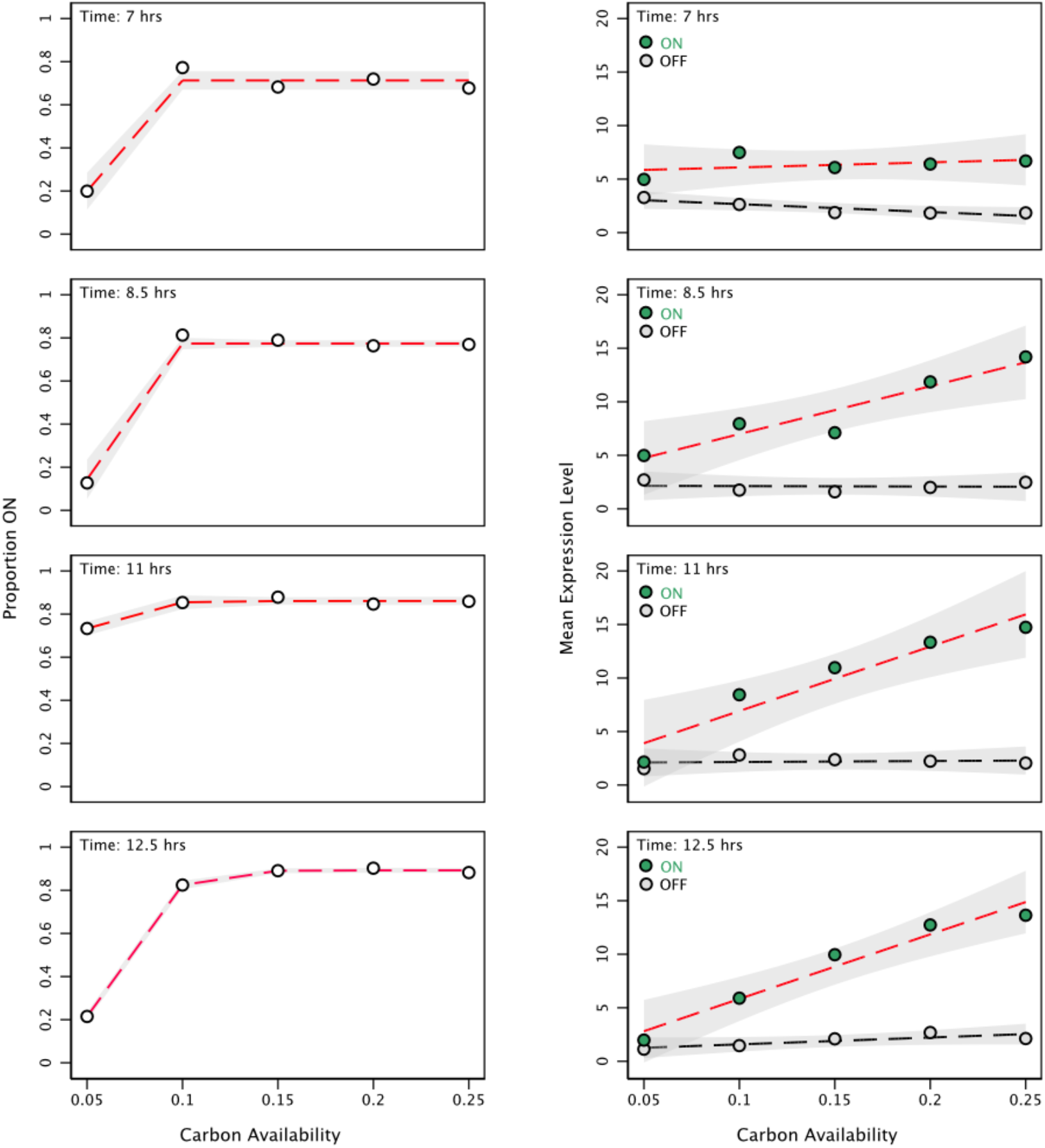
Single cell results are not sensitive to the exact time sampled. In order to test the generality of our results and how sensitive they are to the specific time sampled, we repeated our main experiment with PAO1 pMHLAS, but took samples from five different time points instead of just entry into stationary phase. We observed the same bimodal response and shifts in proportion responding and level of response across all timepoints, concluding that our results are not sensitive to the exact measurement time.

**Figure S6.**
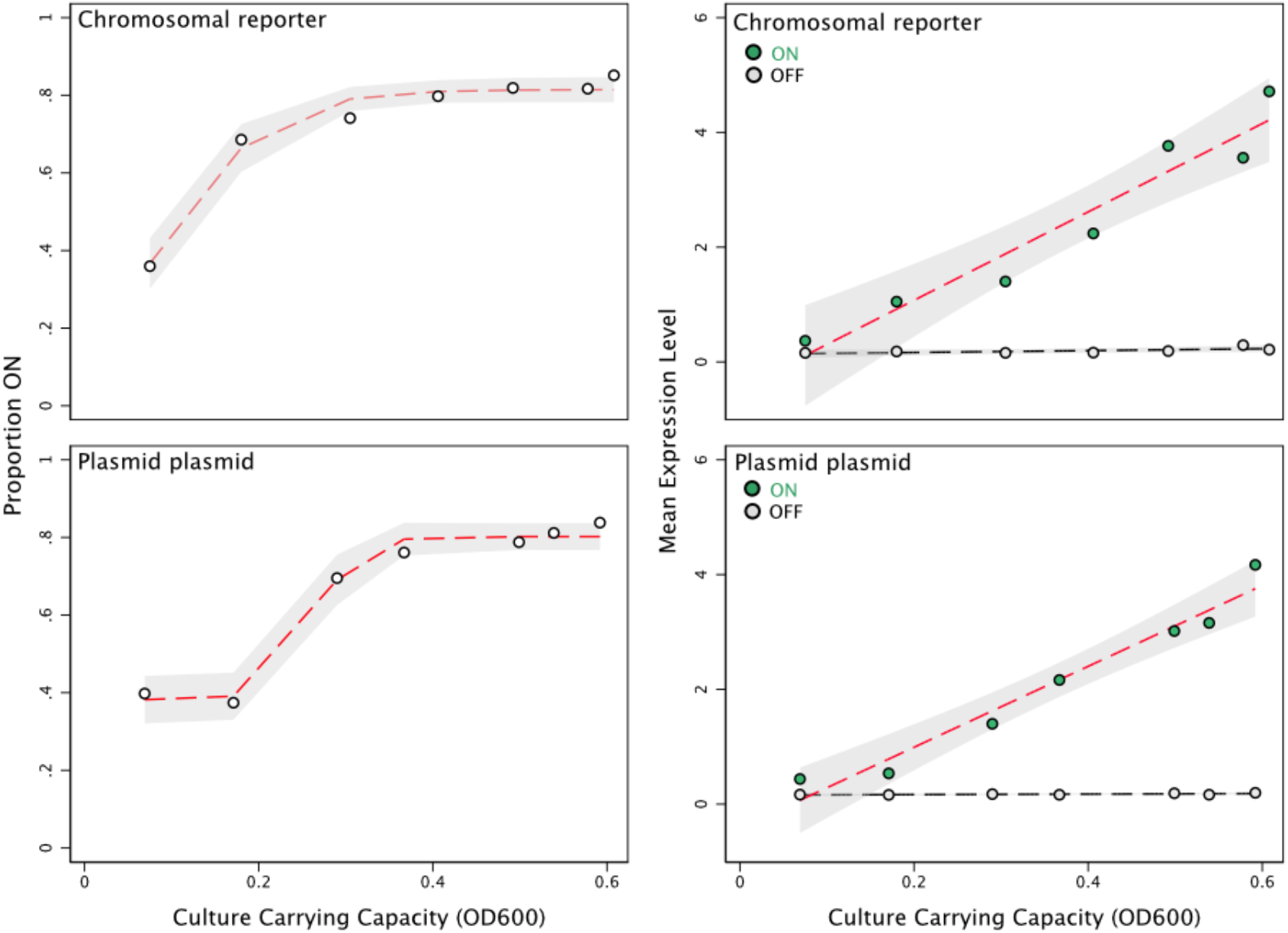
Presence/absence of *Plac::lasR* and plasmid nature of the reporter do not impact graded QS response. The plasmid hosting our reporter, pMHLAS, also contains P*lac::lasR*. While the lac operon is typically repressed, some strains of *Pseudomonas* are capable of leaky, non-induced expression [71]. To confirm our single cell data, we repeated the experiment using a strain with the P*lasB::gfp(ASV)* reporter inserted with the mini-Tn5 method. This strain does not contain P*lac::lasR* and expresses WT levels of *lasR*. We found the same bimodal response to density environments in both strains. This indicates that neither the presence of P*lac::lasR* nor the plasmid-based nature of our reporter are responsible for the graded QS response. This could mean either there is not an appreciable amount of *lasR* being made from the P*lac::lasR* or that the increase of *lasR* does not impact the observed response. These results confirm that the observed response across density is not sensitive to plasmid copy number or the presence of potentially leaky P*lac::lasR* with this specific reporter.

**Figure S7.**
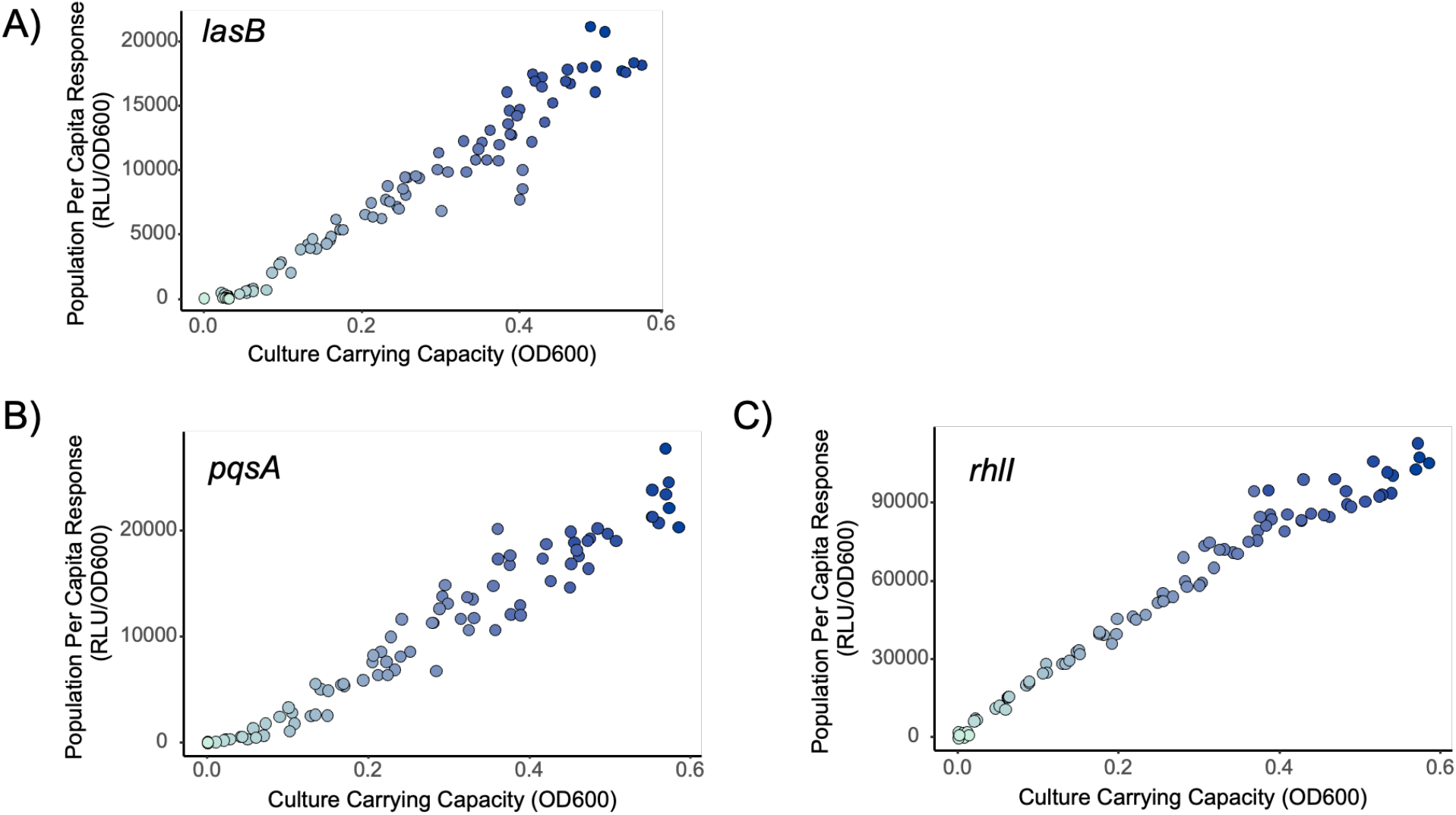
Population response to increasing cell density using chromosomally inserted lux based QS reporters. A) strain P NPAO1 mini-CTX PlasB::lux, B) NPAO1 mini-CTX PpqsA::lux, and C) NPAO1 mini-CTX PrhlI::lux. Bacteria were grown following the same methods outlined in the main text methods and monitored in a plate reader. Per capita expression (RLU/OD_600_) was calculated by taking the maximum expression (RLU) divided by the carrying capacity of the culture (OD_600_). With all three QS reporters, a linear fit model supports the data more than a step-function fit (lasB, AIC linear: 0, AIC step-function: 281.55. pqsA, AIC linear: 1.22, AIC step-function: 283.43. rhlI, AIC linear: 40.68, AIC step-function: 351.86).

**Figure S8.**
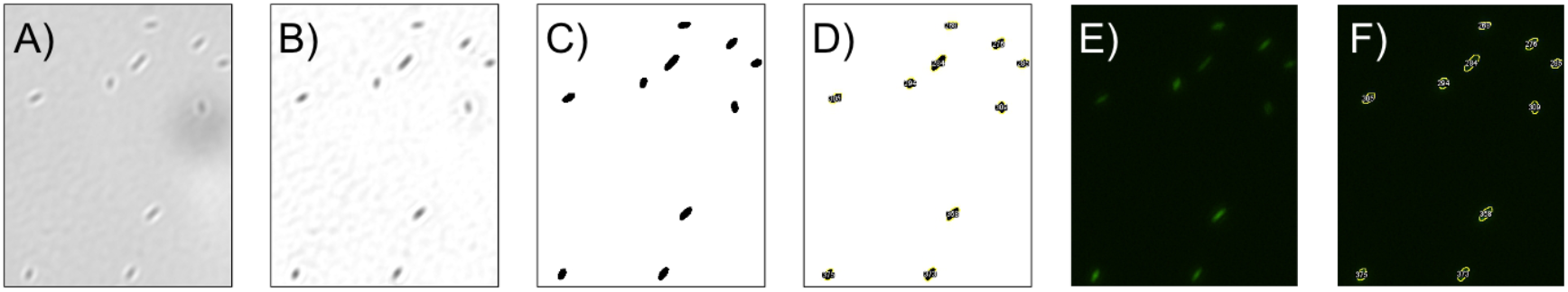
Visual summary of single-cell microscopy analysis pipeline. A) Phase contrast image of individual cells immobilized on 0.01% poly-l-lysine coated glass slides. B) ImageJ’s “background subtraction” command was used to increase contrast between the cells and the background. C) A mask of the phase contrast channel was then created using ImageJ’s default auto threshold, a variation of the IsoData algorithm. D) ImageJ’s “analyze particles” command was then used to identify the features (cells) from the image, this generates a set of regions of interest (ROIs), shown in yellow superimposed. E) Unaltered fluorescence channel image of the same cells. F) ROIs generated from the phase contrast image are then be superimposed onto the unedited fluorescence channel image, either red fluorescence for dead cell identification via propidium iodide or green florescence for the QS reporter, and ImageJ’s “measure” command was used to find the average pixel intensity within each ROI. Any ROIs with red fluorescence were identified as compromised cells and removed from further analysis.

**Table S1.** Strains used in this manuscript.

| Strain/Plasmid | Relevant Characteristics | Figure | Source or Reference |
| --- | --- | --- | --- |
| NPAO1 pMHLAS | QS reporter<br>*PlasB::gfp(ASV)*,<br>*Plac::lasR* | 2, 3, 4, S1, S2,<br>S4, S5 | Hentzer et al., 2002<br>[42] |
| NPAO1 $\Delta lasI \Delta rhII$<br>pMHLAS | Double signal synthase<br>mutant, QS reporter<br>*PlasB::gfp(ASV)* | 2, 3, 4 | This paper |
| PAO1 MH340 mini-Tn5<br>*PlasB::gfp(ASV)* | QS reporter<br>*PlasB::gfp(ASV)*<br>inserted via the mini-Tn5 method | S5 | Kasper Nørskov<br>Kragh |
| NPAO1 mini-CTX *PlasB::lux* | luxCDABE based [72]<br>QS reporter [73]<br>inserted at the *attB* site<br>[74] | S6 | This paper |
| NPAO1 mini-CTX *PpqsA::lux* | luxCDABE based [72]<br>QS reporter [73]<br>inserted at the *attB* site<br>[74] | S6 | This paper |
| NPAO1 mini-CTX *PrhII::lux* | luxCDABE based [72]<br>QS reporter [73]<br>inserted at the *attB* site<br>[74] | S6 | This paper |

### Analysis Supplement

**Figure S9.**
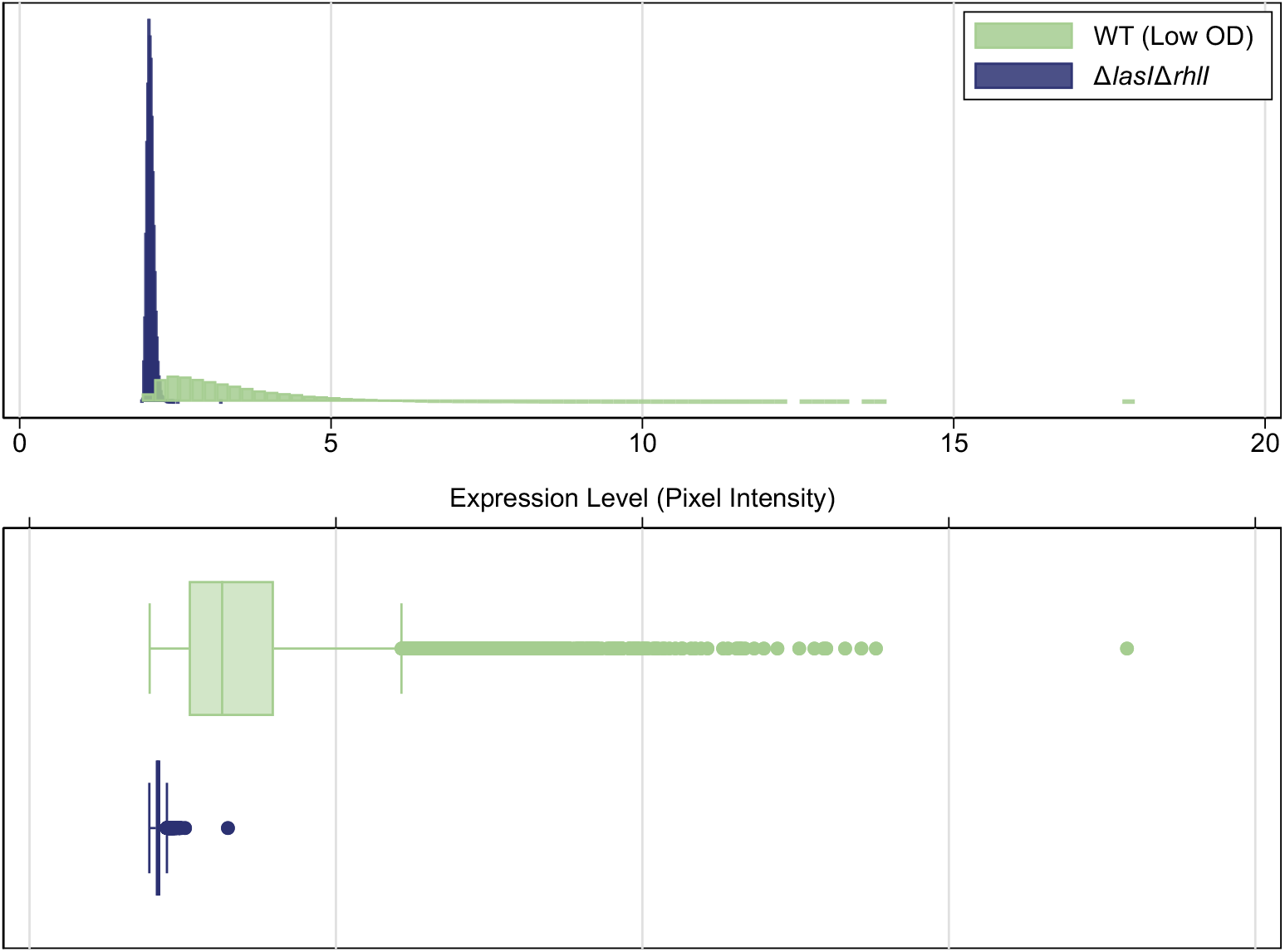
Comparison of pixel intensity distribution for Δ*lasI*Δ*rhlI* and wildtype strains. Figure S1 shows the observed pixel intensity measurements for Δ*lasI*Δ*rhlI* cells in comparison to those for wildtype cells at the lowest measured optical density (OD). As the Δ*lasI*Δ*rhlI* strain is incapable of synthesizing auto-inducers, it should exhibit minimal *lasB* expression. The standard deviation is for expression in the wildtype is1.23. In contrast, pixel intensity for the Δ*lasI*Δ*rhlI* strain is confined to a very narrow range; its standard deviation is 0.06. These results are consistent with a mutant strain that does not express *lasB* suggesting that the mutant’s pixel intensity is near the measurement sensitivity of the experiment. Indeed, the minimum pixel intensity observed in the mutant is 1.955, a value consistent with the overall minimum observed intensity of 1.952. Treating this value as the measurement limit, we subtract it from all measured intensities so that an intensity of 0 corresponds to no expression. The main text figures use *adjusted expression level* to indicate the measured pixel intensity after subtracting this minimum value.

**Figure S10.**
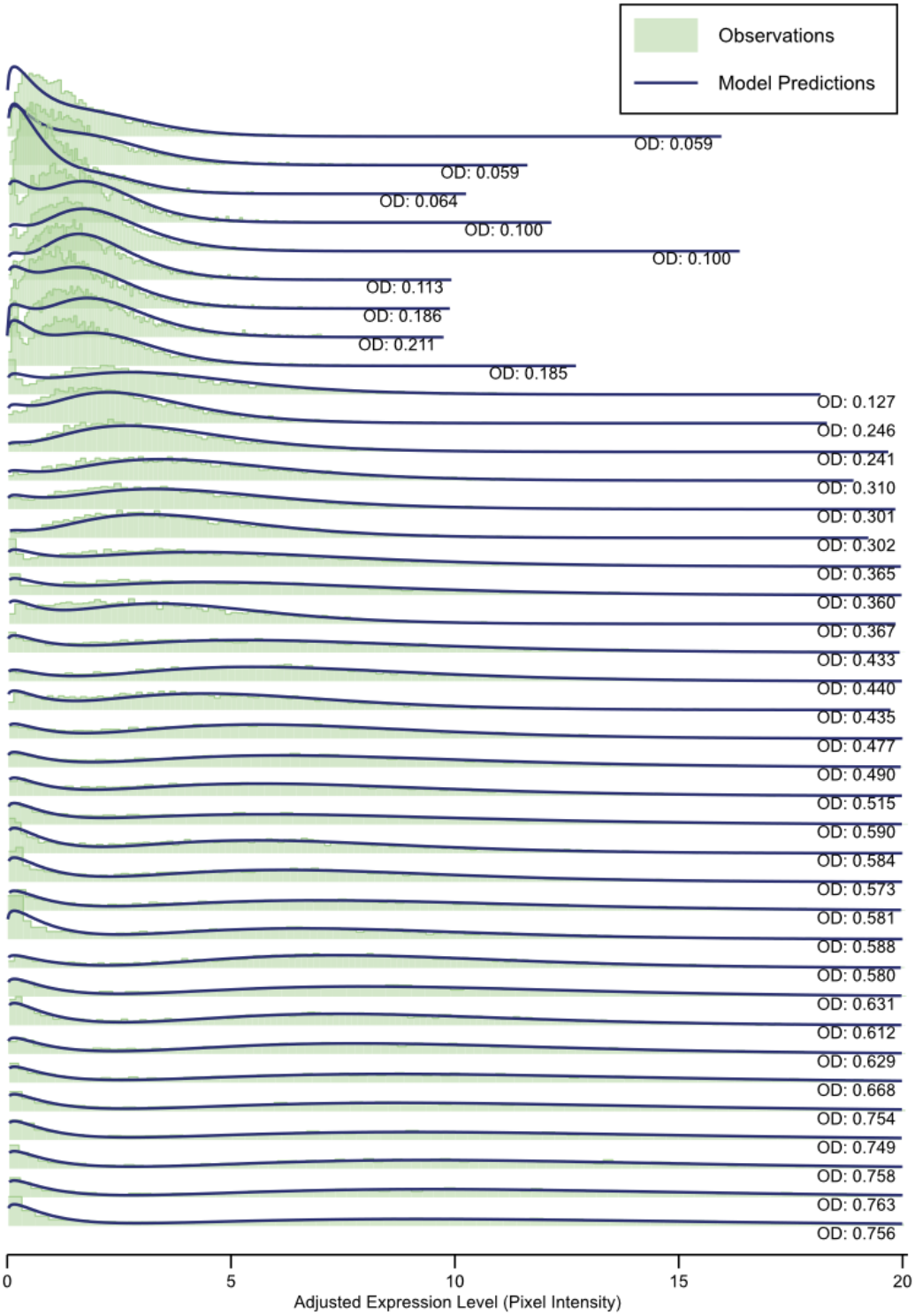
Two-component Gamma mixture model of expression level distributions overlaid on observation histograms. After having identified two sub-populations of cells (Figure S3, Figure S4) we model the observed *lasB* expression levels as a two-component finite mixture model. The sub-populations serve as a latent classes in the model. As gene expression level is non-negative and necessarily right-skewed, we use the Gamma distribution to model expression levels of each class [75,76]. We use expectation-maximization [77,78] to find maximum likelihood estimates for the latent class probabilities and Gamma distribution parameters at each density. Our initial model places no constraints on the parameters other than those required by the model structure (i.e. probabilities between 0 and 1, shape and scale positive). Three parameters from the mixture model characterize the sub-populations of cells: the proportion of cells in the ON sub-population and the mean *lasB* expression levels of the ON and OFF sub-populations. To improve the model’s discriminatory power at low densities, we constrain the OFF sub-population parameters to be those values from the higher densities and update the estimates solely for the ON sub-population. The highest optical density values demonstrate statistically significant bimodality. As expected, the mixture model accurately captures the two sub-populations that make up these distributions. Figure S11 compares these models with the observations.

**Figure S11.**
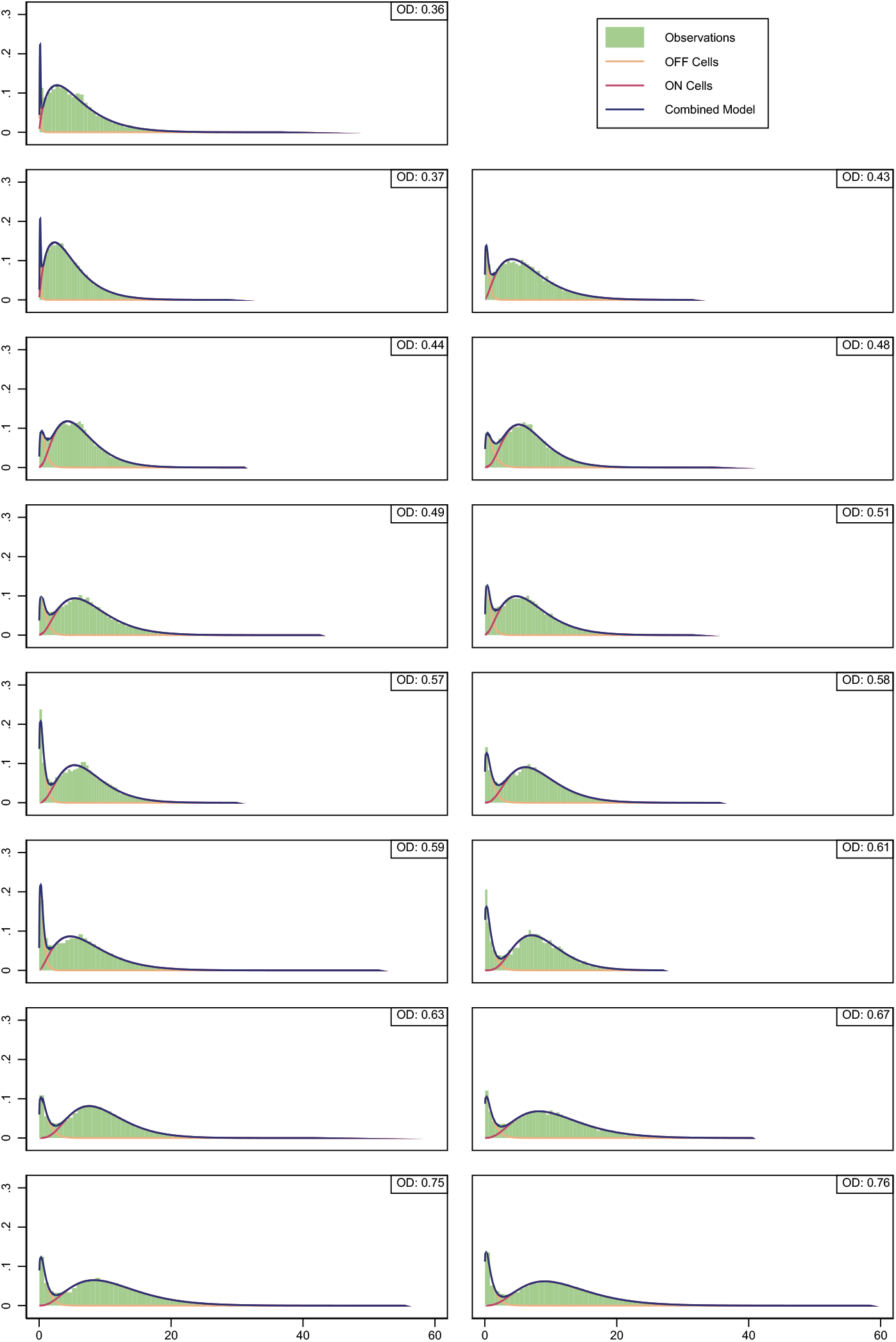
Comparing model predictions with observations for optical density values that demonstration statistically significant bimodality. In addition to the complete model (shown in blue), the plots also show the individual sub-populations of OFF cells (yellow) and ON cells (red). At medium optical density values, the mixture model predictions continue to show close agreement with the observations. Figure S12 highlights this agreement. OD values that don’t demonstrate statistically significant bimodality still show the effects of a mixture of two sub-populations.

**Figure S12.**
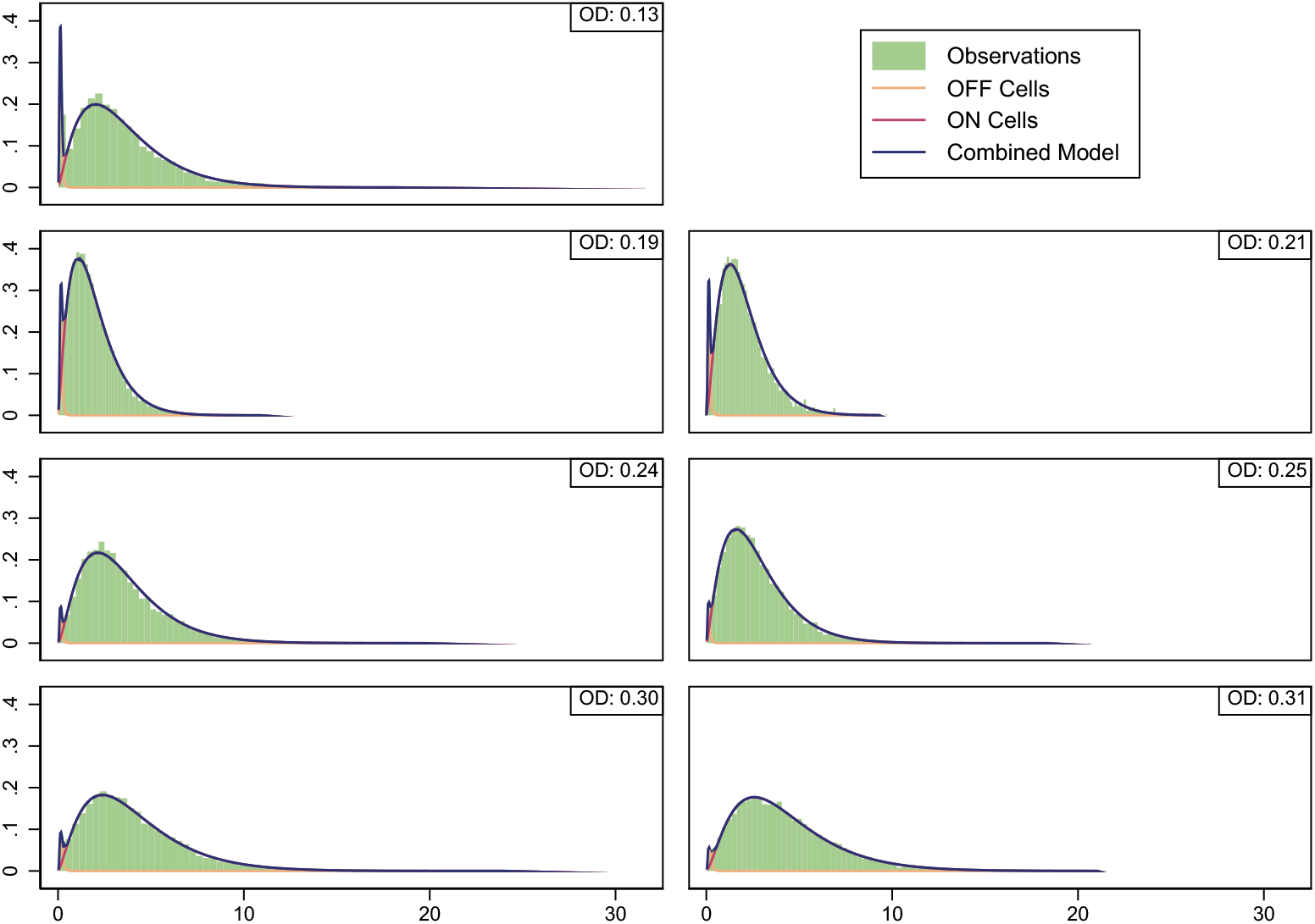
Comparing model predictions at moderate optical densities. As above, the plots show the individual sub-populations of OFF cells (yellow) and ON cells (red). At the lowest optical densities, the benefits of a mixture model are less clear. Figure S13 illustrates the difficulty by comparing the mixture model with a maximum likelihood fit to a single Gamma distribution. As Table S2 indicates, both Akaike Information Criteria (AIC) and Bayesian Information Criteria (BIC) suggest that the mixture model is superior, but the visual evidence is less compelling.

**Figure S13.**
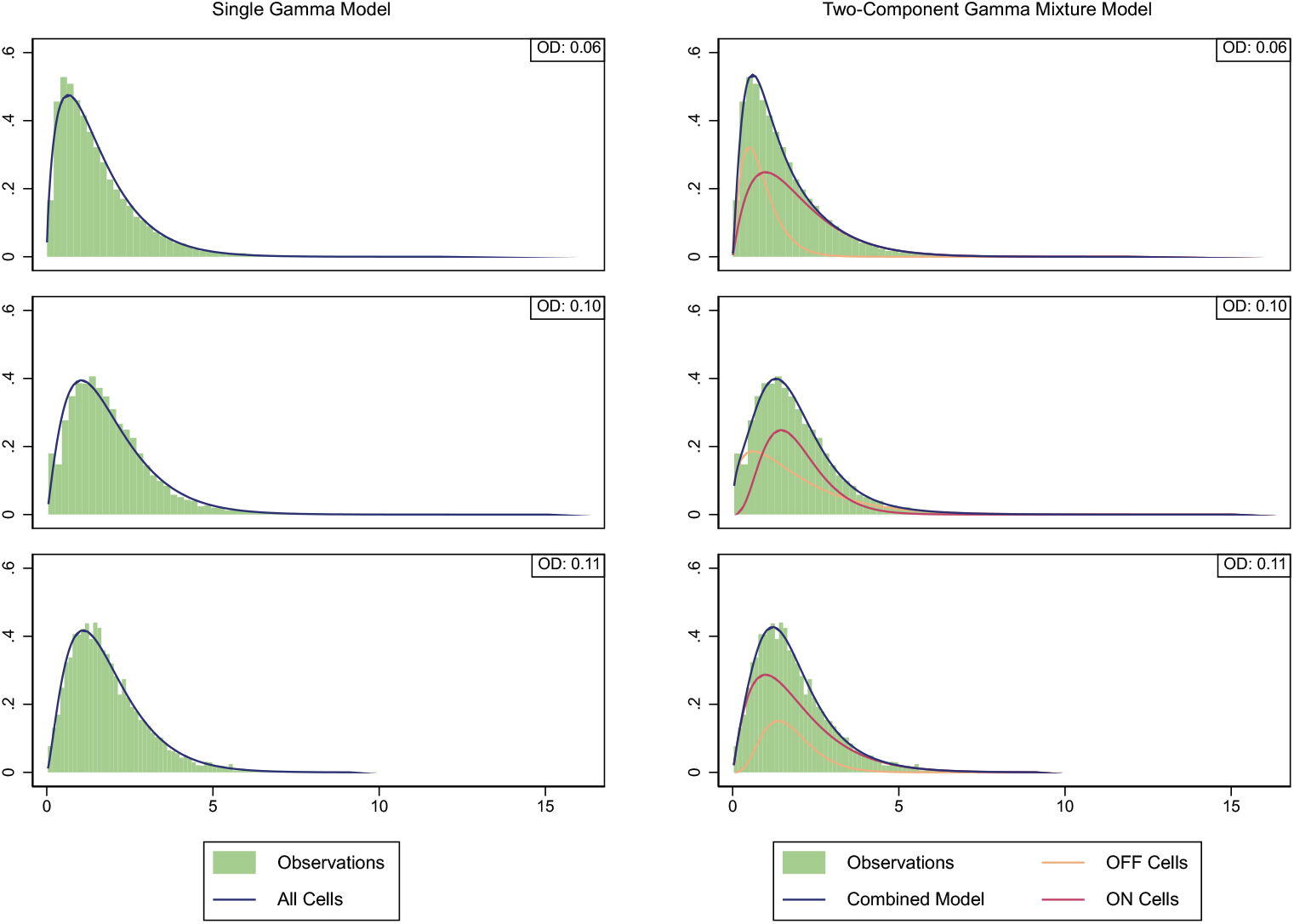
Comparing a model of a single Gamma distribution to the two-component mixture. Plots on the right show a single Gamma distribution with maximum likelihood estimates for shape and scale parameters. Plots on the right show the two-component mixture model. Mixture model plots include individual sub-populations of OFF cells (yellow) and ON cells (red). As the figure and table both show, the observations at optical density 0.11 are especially problematic. At that OD, the statistical mode of *lasB* expression in the ON sub-population is lower than that in the OFF sub-population. As required by the model, however, the mean expression level of the ON sub-population is higher than that of the OFF sub-population.

**Table S2.** Information criteria for single Gamma (subscript 1) and two-component Gamma mixture (subscript 2) models at low optical densities. The Δ columns show change in associated information criteria in two-component model. Final two columns are likelihood ratio and p-value.

| OD | AIC<sub>1</sub> | AIC<sub>2</sub> | ΔAIC | BIC<sub>1</sub> | BIC<sub>2</sub> | ΔBIC | LR $\chi^2(3)$ | p-value |
| --- | --- | --- | --- | --- | --- | --- | --- | --- |
| 0.06 | 55741 | 55472 | -268 | 55757 | 55512 | -245 | 274 | 0 |
| 0.10 | 33499 | 33289 | -210 | 33514 | 33326 | -188 | 216 | 0 |
| 0.11 | 20994 | 20972 | -22 | 21008 | 21007 | -1 | 28 | $3.5 \times 10^{-6}$ |

